# Enhanced Antibody Response to the Conformational Non-RBD Region *via* DNA Prime-Protein Boost Elicits Broad Cross-Neutralization Against SARS-CoV-2 Variants

**DOI:** 10.1101/2024.02.04.578544

**Authors:** Yun-Fei Ma, Kun Chen, Bowen Xie, Jiayi Zhu, Xuan He, Chunying Chen, Yuhe Renee Yang, Ye Liu

## Abstract

Preventing immune escape of SARS-CoV-2 variants is crucial in vaccine development to ensure broad protection against the virus. Conformational epitopes beyond the RBD region are vital components of the spike protein but have received limited attention in the development of broadly protective SARS-CoV-2 vaccines. In this study, we used a DNA prime-protein boost regimen to evaluate the broad cross-neutralization potential of immune response targeting conformational non-RBD region against SARS-CoV-2 viruses in mice. Mice with enhanced antibody responses targeting conformational non-RBD region show better performance in cross-neutralization against the Wuhan-01, Delta, and Omicron subvariants. *Via* analyzing the distribution of conformational epitopes, and quantifying epitope-specific binding antibodies, we verified a positive correlation between the proportion of binding antibodies against the N-terminal domain (NTD) supersite (a conformational non-RBD epitope) and SARS-CoV-2 neutralization potency. The current work highlights the importance of high ratio of conformational non-RBD-specific binding antibodies in mediating viral cross-neutralization and provides new insight into overcoming the immune escape of SARS-CoV-2 variants.

## Introduction

SARS-CoV-2 virus, responsible for COVID-19, continuously evolves, producing new variants that challenge global health. These variants, with mutations in the spike protein, often exhibit increased transmissibility and immune escape capabilities [1,2]. Vaccines remain the most effective and economical method for preventing COVID-19. However, the continuous emergence of new variants and immune escape of the SARS-CoV-2 viruses makes the development of broadly protective vaccines increasingly urgent.

The spike (S) protein is the primary antigen for developing SARS-CoV-2 vaccines. The S protein comprises the receptor-binding domain (RBD) and non-RBD regions [3]. Given its critical role in mediating viral entry, the RBD has been a predominant target in therapeutic and vaccine research [4–6]. However, the high frequency of mutations in the RBD region reduces its significance in broad-spectrum SARS-CoV-2 vaccine research [7–10]. In contrast, the non-RBD region mutates at a lower and slower frequency [11–13], suggesting a more promising potential in limiting the immune escape from viral variants. The N-terminal domain (NTD) is a critical component of the non-RBD region of the spike protein. Some neutralizing antibodies (nAbs) targeting the NTD have been identified, exhibiting potent and broad neutralization against diverse SARS-CoV-2 variants [14–16]. Furthermore, evidence suggested that incorporating the NTD into vaccine designs enhanced their capacity for broad cross-neutralization [17–19]. However, non-RBD-based broad-spectrum vaccine research remains limited.

Preserving the spatial conformation of antigens is critical for effectively inducing neutralizing antibody responses [20–22]. For SARS-CoV-2 vaccines, the RBD as core antigen can preserve a satisfactory spatial conformation [21,23,24] to induce immune response. Nevertheless, how to preserve the spatial conformation of epitopes in non-RBD region still suffers a big challenge [25,26]. The non-RBD region is divided into two discontinuous fragments by the RBD domain [27]. If non-RBD antigens are harvested *via* fragment-fusion or fragment-separated expression *in vitro*, the spatial conformation of epitopes in non-RBD region would very possibly undergo a significant change [28]. Therefore, it is very challenging to obtain conformational non-RBD antigen by the traditional protein expression strategy. To understand the roles of conformational non-RBD in enhancing the broad cross-neutralization of vaccines, we would like to indirectly compare the immune responses between the full-length S protein (including conformational non-RBD antigen) and the RBD antigen.

As the DNA prime enhanced T helper and B cell responses and induced strong antibody responses against various pathogens [29–33], we chose the DNA prime strategy. In the current study, we used the ‘DNA prime-protein boost’ strategy to immunize conformational non-RBD region in the mouse model. We found that a high ratio of binding antibodies targeting conformational non-RBD region can significantly enhance cross-neutralization against SARS-CoV-2 viruses. Furthermore, by a combined analysis of intelligentized epitope identification and electron microscopy (EM) techniques, we identified NTD supersite as a critical conformational epitope in the non-RBD region contributing to increased SARS-CoV-2 cross-neutralization. This finding shed light on the underlying mechanism of conformational non-RBD epitopes in mediating cross-neutralizing against SARS-CoV-2 viruses.

## Materials and methods

### Vaccine design

The Spike protein of SARS-CoV-2, spanning positions 21,563–25,384 in the viral genome, was codon-optimized for expression in 293 cell lines. Similarly, the RBD protein, encompassing positions 222,517–23,185 in the SARS-CoV-2 isolate Wuhan-Hu-1 (GenBank accession number: MN908947), underwent codon optimization for expression in CHO-S cell lines. Codon-optimized sequences encoding the full-length spike (S) or RBD protein of SARS-CoV-2 were inserted into PVAX-1 expression plasmids, serving as DNA vaccines.

### Protein expression and purification

A plasmid encoding the SARS-CoV2 S-2 S protein and RBD protein, incorporating an FC-tag for purification, was codon-optimized and synthesized by GenScript. Subsequently, these constructs were cloned into pCDNA2004. The chosen expression platforms were 293F and CHO-S cells. Transient transfection of cells was performed utilizing Hieff TransTM Liposomal Transfection Reagent (Yeasen Biotechnology) following the manufacturer’s instructions. Cultures were maintained for 6 days at 37 °C and 5% CO_2_. Following transfection, the culture supernatant was collected and centrifuged for 5 min at 300 × g to eliminate cells and cellular debris. The resulting supernatant underwent sterile filtration using a 0.22 µm vacuum filter and was subsequently stored at 4 °C until needed. Purification of the FC-tagged SARS-CoV-2 Spike protein and RBD protein was carried out employing a purification protocol employing 5 ml Protein At Beads 4FF (Smart-Lifesciences). FC tags on the S and RBD proteins were removed through treatment with Factor Xa Proteinase (Promega).

### Immunizations

Female BALB/c mice, aged six to eight weeks, were sourced from the Institute of Medical Biology, Chinese Academy of Medical Sciences (CAMS), and were maintained in specific pathogen-free (SPF) conditions at the Central Animal Care Services of Institute of Medical Biology, CAMS. All mouse experiments were ethically approved by the animal ethics committee of the Institute of Medical Biology, CAMS, with the assigned review number DWSP202008021. Experimental and control animals were co-housed, and the research staff received training in animal care and handling. Approval for animal experiments was obtained from the Yunnan Provincial Experimental Animal Management Association.

The mice were randomly divided into five groups (n = 10) as follows: (1) 35 μg aluminum adjuvant and 10 μg Spike protein for three doses (Alu+S protein group); (2) 35 μg aluminum adjuvant and 10 μg RBD for three doses (Alu+RBD group); (3) 15 μg DNA vaccine for one dose, 35 μg aluminum adjuvant and 10 μg Spike protein for two-dose (DNA prime-Alu+S protein boost group); (4) 15 μg DNA vaccine for one dose, 35 μg aluminum adjuvant, and 10 μg RBD for two doses (DNA prime-Alu+RBD protein boost group); (5) 0.9 % saline (Control group).

On day 0, mice received DNA vaccine or saline (Control group) through intramuscular injection. Protein vaccines or saline (Control group) were administered *via* intramuscular injection on day 14 and day 28. Two weeks after the final immunization, mice were euthanized to collect spleens and serum samples.

### Antigen-specific binding antibody assay

The analysis of Spike and RBD antibody responses was conducted using enzyme-linked immunosorbent assay (ELISA). Spike/RBD-specific IgG, IgG1, IgG2a, IgG2b, and IgG3 in mice serum were quantified by ELISA. Specifically, 96-well plates were coated with Spike or RBD antigen (0.25 μg per well) and left overnight at 4 °C. Subsequently, these plates were blocked with 2% BSA for 2 h at 37 °C. After blocking, the plates were washed three times with PBST (Phosphate buffer saline (PBS) with 5‰ Tween 20), and 100 μl mice serum was added per well. Incubation was carried out at 37 °C for 1 h. Following incubation, the plates were washed three times with PBST. Each well of 96-well plate was then treated with 100 μl HRP-labeled goat anti-mouse IgG, IgG1, IgG2a, IgG2b, IgG3, IgM binding antibody at 37 °C for 1 h. Following another three washes with PBST, 50 μl 3,3′,5,5′-tetramethylbenzidine (TMB; Sigma) was added to each well. The reaction was stopped with 50 μl 0.05% H_2_SO_4_, and the absorbance values at 450 nm and 630 nm wavelengths were determined using an ELISA plate reader (Tecan, San Jose, CA).

### Live virus neutralization assay

Neutralizing antibody detection was conducted using a live virus infection platform. Three SARS-CoV-2 live virus types (Wuhan-Hu-1, B.1.1.7, and B.1.617.2) were obtained from the Institute of Medical Biology, Chinese Academy of Medical Sciences (CAMS). Serum serial dilutions were incubated with 2000 PFU of an early SARS-CoV-2 isolate in 100 µl OptiPro medium supplemented with 1x GlutaMAX (both Gibco) for 1 h at 37 °C. The mixture was then added onto a confluent monolayer of Vero E6 cells (seeded the day before at 10^4^ cells per well in a 96-well plate). After 1 h, the mixture was removed, and 100 µl OptiPro medium supplemented with 1x GlutaMAX (both Gibco) was added. After 24 h of incubation at 37 °C and 5% CO_2_, cells were fixed with 100 µl 4% paraformaldehyde for 20 min at room temperature (RT) and permeabilized with 100 µl 0.5% Triton X-100 in PBS for 15 min at RT. Following a blocking step with 100 µl 5% skimmed milk in PBS for 1 h at RT, cells were stained with purified immunoglobulins from a SARS-CoV-2 convalescent patient in 2% skimmed milk for 1 h at 4 °C. After three washing steps with 200 µl PBS, 100 µl of anti-human IgG FITC (1:200, Jackson Immunoresearch) diluted in 2% skimmed milk were added and incubated for 1 h at 4 °C in the dark. After another three washing steps with 200 µl PBS, plaques were counted using an ELISPOT reader and analyzed with CTL Immunospot software (Cellular Technology limited BioSpot). Infected wells without serum were used as a reference to determine the 75% plaque reduction neutralization titer (PRNT75).

### Blocking NTD-specific antibodies

SARS-CoV-2 pseudotyped virus (Wuhan-Hu-1) was obtained from the Institute of Medical Biology, Chinese Academy of Medical Sciences (CAMS). For the blocking experiment, 100 µl of DMEM supplemented with 10% FBS was added to each well of a 96-well round-bottom plate. For the low-concentration NTD protein blockade (0.04 μg/μL), 44 µl of DMEM (+10% FBS), 3 µl of serum, and 3 µl of NTD protein (2.05 mg/mL, AINOD) were added to column 1. For the high-concentration NTD protein blockade (0.08 μg/μL), 41 µl of DMEM (+10% FBS), 3 µl of serum, and 6 µl of NTD protein (2.05 mg/mL, AINOD) were added to column 1. For the control group, 47 µl of DMEM (+10% FBS) and 3 µl of serum were added to column 1. Place the plate into a 37 °C and 5% CO_2_ incubator for 1 hour. Transfer 50 µl serum from column 1 into column 2 and mix thoroughly. Similarly transfer 50 µl from column 2 to column 3, column 3 to column 4, column 4 to column 5, and column 5 to column 6. After mixing in column 6, discard 50 µl of serum dilution. Transfer 50 µl of pseudotyped viruses (Wuhan-Hu-1) into the serum dilution in the round bottom 96-well plate (except for the No virus control wells)and mix well. Place the plate into a 37 °C and 5% CO_2_ incubator for 1 hour. Resuspend and count 293T-hACE2 cells in DMEM (+ 10% FBS). Dilute the cell suspension to 2 x 10^5^ cells/mL and transfer 100 µl into each well of 96-well flat bottom plate using a multichannel pipette. Put the 96-well plate into an incubator set at 37 °C and 5% CO_2_ incubator for 48 h. Remove the medium from the wells. Add 20 µl/well of 1x luciferase cell culture lysis reagent (Promega). Add 100 µl of Luciferase Assay reagent (Promega) to each well. Measure the light produced (BioTek: CYTATION). The serum neutralization titers and inhibitionand of pseudotyped virus were analyzed using Prism 9 (GraphPad). Statistical analyses were performed using a two-tailed Student’s t-test (GraphPad Prism 9).

### Serum IgG isolation and digestion

For IgG isolation from the serum, the following procedure was employed. Heat 500 μl of mouse serum at 56 °Cand then incubate the heated serum with 1 ml of washed Protein-G resin (GenScript L00209) overnight at 4°C. Wash the resin complex three times with PBS through centrifugation. Elute the complex with 5 ml of 0.1 M glycine buffer at pH 2.0. Immediately neutralize the eluate with 4 ml of 1 M Tris-HCl (pH 8.0) and filter the neutralized eluate using a 0.2 μm pore filter. Buffer exchanged the elution to PBS using 30-kDa cut-off Amicon ultrafiltration units. For Fab preparation: incubate 500 μg of concentrated polyclonal IgG samples with papain (Sigma Aldrich P3125) in digestion buffer (100 mM Tris, 2 mM EDTA, 10 mM L-Cysteine, pH 7.4) for approximately 4 h in a 37°C incubator. Add iodoacetamide (Sigma Aldrich I1149-25G) to quench the digest. Then concentrate the digestion and buffer-exchanged to TBS using 10-kDa cut-off Amicon ultrafiltration units. Assess the digestion for purity using SDS-polyacrylamide gel electrophoresis.

### Identifying and partitioning epitopes of neutralizing antibody

We gathered information on antibodies targeting and neutralizing the spike protein of SARS-CoV-2 from the Coronavirus Antibody Database (CoV-AbDab: http://opig.stats.ox.ac.uk/webapps/covabdab/) through filtering. The structures of neutralizing antibodies bound to the SARS-CoV-2 spike protein were obtained from the Protein Data Bank (PDB: https://www.rcsb.org/).

To efficiently and accurately classify hundreds of epitope residues targeted by various neutralizing antibodies, we developed a method composed of three iterative cycles. In the initial cycle, we start with the epitope residues of a randomly selected neutralizing antibody. These epitope residues are merged with the epitope residues of other antibodies if the overlapping ratio between the merged set and the epitope residues of another antibody is no less than 0.9. If no epitope residues meet the merger criteria, the process concludes. We then restart again with the epitope residues of a randomly selected antibody from the remaining antibodies until all antibodies are processed. After completing the first cycle, the output becomes the input for the second iterative process, repeated three times. In this process, we reduce the overlapping ratio to 0.8. Finally, the output from the second iterative process serves as input for the last cycle, repeated only two times, with the overlapping ratio reduced to 0.6. The number of final epitope regions decreases with the reduction of the overlapping ratio, and reducing the number of iterative cycles increases the number of final epitope regions.

The epitope site of a neutralizing antibody is defined as the antigen residues contacted and buried by the neutralizing antibody. A contacted epitope residue refers to an antigen residue that contacts the antibody within a distance of 5 Å in the spike-antibody complex, while the buried epitope is defined as an antigen residue with a buried surface area (BSA) greater than 0 Å^2^. Hydrogen bonds between the antigen and antibody were calculated using PyMOL.

In summary, our method enables the rapid and accurate classification of hundreds of epitope residues targeted by different neutralizing antibodies. The efficiency of our method stems from the iterative cycle that merges epitope residues, reducing the time and complexity of manual residue comparison. The accuracy results from the thoroughness of the iterative cycle, ensuring all epitope residues are considered.

### Preparation of Fab-spike complexes for ns-EMPEM

Fab-spike complexes were generated through the incubation of 15 μg of spike protein with 500 μg of polyclonal Fabs overnight at 4 °C. Subsequently, the complexes were subjected to purification using a Superdex 200 Increase 10/300 column with UV absorbance at 215 nm on Akta Pure system (cytiva). The purification was performed in tris-buffered saline (TBS) buffer. Fractions containing the spike-Fab complexes were collected and concentrated using 10-kDa cutoff Amicon ultrafiltration units. The concentrated complexes were then immediately utilized for preparing ns-EMPEM grids.

### Ns-EMPEM sample preparation and data collection

In the ns-EMPEM analysis, antibodies were examined using spike-Fab complexes. The final concentration of the collected fraction was approximately 0.015 µg/µL. Complexes were diluted to 0.015 µg/mL in 1× Tris-buffered saline and 3.5µl applied to a 200 mesh Cu grid (Zhongjingkeyi (Beijing) Films Technology Co., Ltd) blotted with filter paper, and stained with 0.7% uranyl formate (Electron Microscopy Sciences. 22400. USA). Micrographs were collected ∼100-120 on a JEOL JEM-2100F high-resolution transmission Electron microscope operating at 200LkV with a camera at ×30,000 magnification using the serial EM automated image collection software. EM-map reconstruction was performed using Relion 3.1.3, and the maps were aligned and displayed using Chimera. Ns-EM density maps were deposited under EMD-39176, EMD-39179, EMD-39181, EMD-39183.

## Results

### DNA prime-protein boost elicits higher levels of immune response Against SARS-CoV-2 virus

To compare the immune response and neutralizing efficacy of the ‘DNA prime-protein boost’ strategy versus traditional protein vaccination, BALB/c mice (n = 10) were immunized intramuscularly as outlined in Figure 1 (A and E). Both the RBD subunit and full-length spike antigens were assessed. Serum samples were collected two weeks after the final immunization, and RBD- and spike-specific antibody responses were quantified via enzyme-linked immunosorbent assay (ELISA). Antigen-specific IgG responses in the ‘DNA prime-protein boost’ groups were 2.82-fold (*P* = 0.02) and 1.67-fold (*P* = 0.0077) higher than those in the RBD and spike protein-only groups, respectively (Figure 1 (B, F)). Additionally, antibody subtypes against RBD and spike were measured by ELISA, revealing that the ‘DNA prime-protein boost’ strategy significantly induced higher levels of IgG2a and IgG2b compared to protein-only immunization for both antigens (Figure 1 (C, G)).

**Figure 1.**
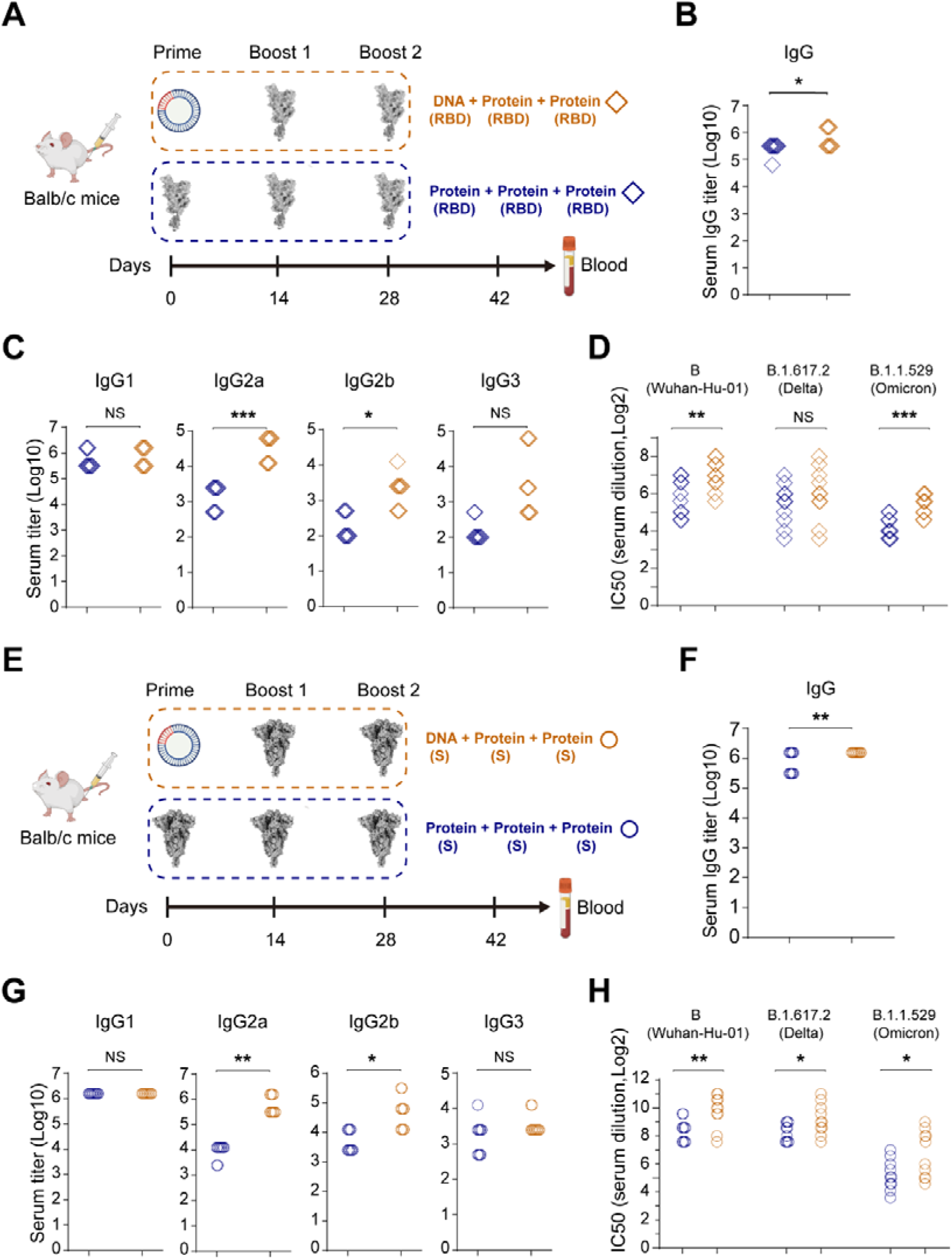
Humoral responses of ‘DNA prime-protein boost’ and protein-only immunization strategies. (A) Schematic diagram of RBD antigen immunization. (B) RBD-specific IgG was assessed by ELISA. (C) RBD-specific IgG subtypes were assessed by ELISA. (D) Live virus neutralization titers were determined by *in vitro* neutralization assays. (E) Schematic diagram of Spike protein immunization. (F) S-specific IgG was assessed by ELISA. (G) S-specific IgG subtypes were assessed by ELISA. (H) Live virus neutralization titers were determined by *in vitro* neutralization assays. The orange and blue rhombuses represent the RBD ‘DNA prime-protein boost’ group and the RBD protein group, respectively (B-D). The orange and blue circles represent the S ‘DNA prime-protein boost’ group and the S protein group, respectively (F-H). Each rhombus (B-D) or circle (F-H) represents one sample, n = 10. \**P* < 0.05, \*\**P* < 0.01, \*\*\**P* < 0.001 (two-tailed Student t-test). NS, not significant.

The neutralizing antibody responses of the ‘DNA prime-protein boost’ and protein-only strategies were further evaluated using live virus neutralization tests. For the RBD antigen, mice immunized with the ‘DNA prime-protein boost’ strategy exhibited 2.16-, 1.52-, and 3.28-fold higher neutralizing antibody responses against the Wuhan-Hu-01, B.1.617.2 (Delta), and B.1.1.529 (Omicron) strains, respectively (Figure 1 (D)). Similarly, in the spike antigen groups, the‘DNA prime-protein boost’ strategy elicited serum neutralizing antibody titers averaging 2.92-, 2.28-, and 3.78-fold higher than those observed with spike protein-only immunization against the same viral strains (Figure 1 (H)).

These findings indicate that the ‘DNA prime-protein boost’ vaccination induces significantly stronger binding and cross-neutralizing antibody responses compared to protein-alone vaccination. Collectively, the ‘DNA prime-protein boost’ strategy markedly enhances the vaccine’s efficacy in eliciting robust humoral responses in the host.

### Boosting conformational non-RBD epitopes to improve SARS-CoV-2 cross-neutralizing response

Before designing the immunization strategy, we investigated whether *in vitro*-expressed non-RBD and RBD antigens were suitable for directly boosting corresponding epitopes. We calculated the RMSD value of non-RBD (w/o linker) and non-RBD (with linker) antigen for 18 SARS-CoV-2 strains (Supplemental Figure 1 (A) and Supplemental Table 1). The RMSD for non-RBD _(w/o linker)_ ranged from 4.0 to 13.1 Å, and for non-RBD _(with linker)_ from 3.9 to14.3 Å, both of which exceed 3 Å (Figure 2 (A and B), Supplemental Figure 1 and 2). These results indicate that *in vitro*-expressed non-RBD antigen hardly preserves its spatial conformation. In contrast, the RMSD value for RBD antigen ranged from 0.5 to 1.6 Å (< 3 Å, Figure 2 (A-B), Supplemental Figure 1 and 2), suggesting that *in vitro*-expressed RBD antigen can maintain its spatial conformation.

**Figure 2.**
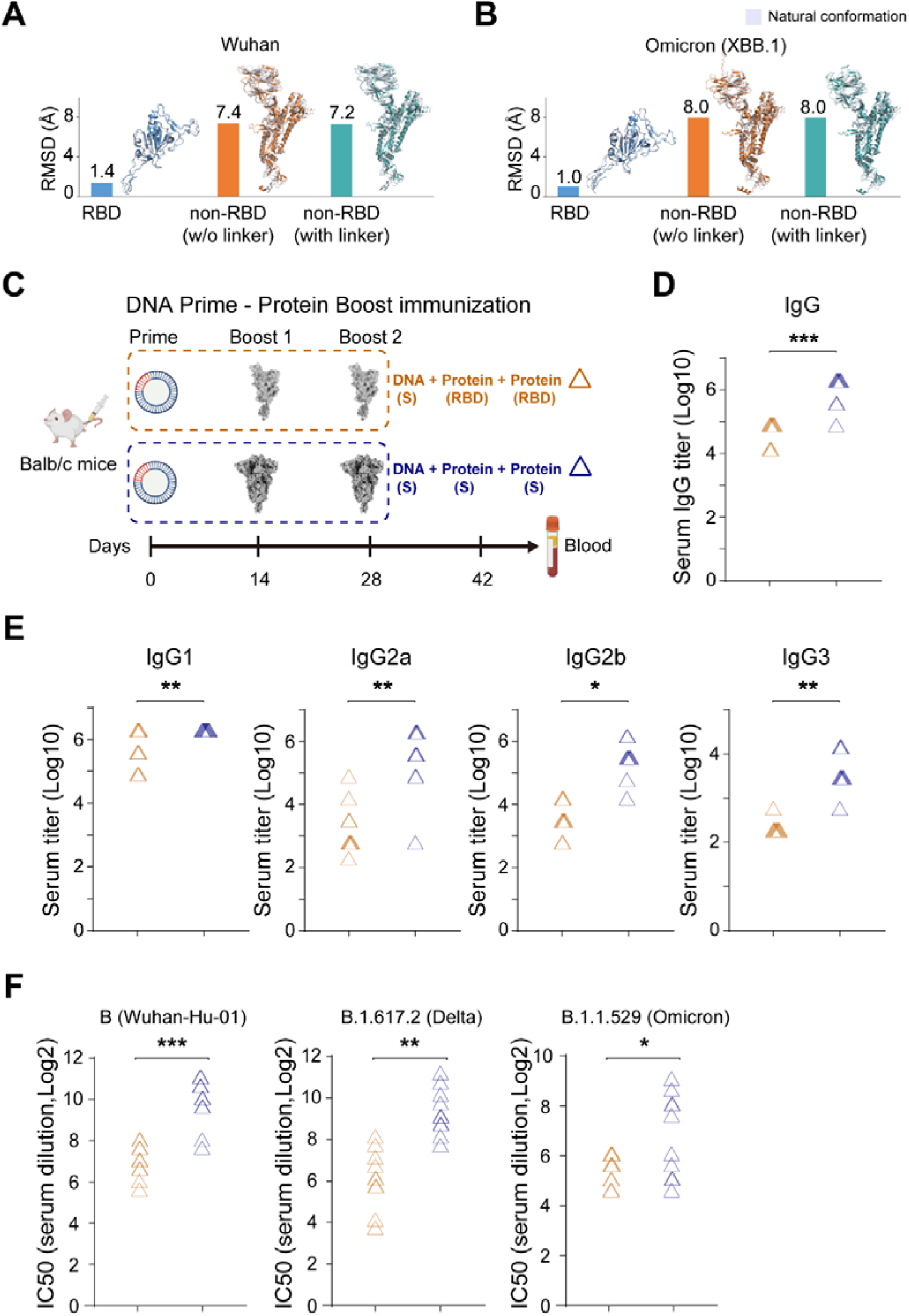
Humoral responses of conformational non-RBD and RBD epitopes. (A-B) Structure analysis of conformational non-RBD and RBD subunits. (C) Schematic diagram of immunization. (D) Spike-specific IgG was assessed by ELISA. (E) Spike-specific IgG subtypes were assessed by ELISA. (F) Live virus neutralization titers were determined by *in vitro* neutralization assays. The orange and blue triangles represent the S_DNA_-RBD-RBD group and the S_DNA_-S-S group, respectively (D-F). Each triangle (D-F) represents one sample, n = 10. \**P* < 0.05, \*\**P* < 0.01, \*\*\**P* < 0.001 (two-tailed Student t-test). NS, not significant.

Subsequently, we employed the ‘DNA prime-protein boost’ strategy to assess the role of boosting conformational non-RBD epitopes in cross-neutralizing responses against SARS-CoV-2 viruses. Mice (n = 10) were boosted with either 10 μg RBD protein (conformational non-RBD epitopes were not boosted) or 10 μg full-length S protein (conformational non-RBD epitopes were boosted) (Figure 2(C)). During the prime stage, all mice received a single dose of DNA vaccine (15 μg) expressing the full-length S protein. Serum samples were collected two weeks after the final vaccination to evaluate humoral responses.

The IgG titer was increased by 23.47-fold in mice receiving a full-length S protein boost (S_DNA_-S-S), in comparison to those receiving an RBD boost (S_DNA_-RBD-RBD) (Figure 2(D)). IgG1, IgG2a, IgG2b and IgG3 response in full-length S protein-boost group was 1.11-fold, 87.47-fold, 87.47-fold and 26.32-fold higher than those in RBD-boost group (*P* < 0.05, Figure 2 (E)).

Of note, the live virus neutralization assay revealed that cross-neutralizing responses were significantly enhanced when conformational non-RBD epitopes were boosted *via* injection of the full-length S protein, compared to boosting RBD epitopes alone. The neutralizing antibody responses of mice boosted with full-length S protein were 7.69-, 8.24- and 4.02-fold higher than those boosted with RBD against the Wuhan-01, B.1.617.2 and B.1.1.529 strains, respectively (Figure 2 (F)). These findings demonstrated that boosting conformational non-RBD epitopes can significantly improve antibody response and cross-neutralization against SARS-CoV-2 viruses.

### Analyzing the distribution of neutralizing epitopes in conformational non-RBD and RBD regions

Clarifying the spatial distribution of primary antigenic epitopes on the S protein holds critical significance in unveiling the mechanisms underpinning cross-neutralization against SARS-CoV-2 variants, particularly those mediated by the conformational non-RBD region. To investigate the distribution pattern of non-RBD and RBD neutralizing epitopes in spike protein, we conducted a comprehensive analysis of 2,397 neutralizing antibodies (nAb) sequences and structures from the coronavirus antibody database (CoV-AbDab) [34] (Figure 3 (A)). After the filtration based on the standard of the immune model, antibody function, structural accessibility, and variant analysis, we picked out 78 entries with crystal structures of spike antigen-neutralizing antibody complexes (S-nAbs) in the protein data bank (PDB) [35] for the epitope analysis.

**Figure 3.**
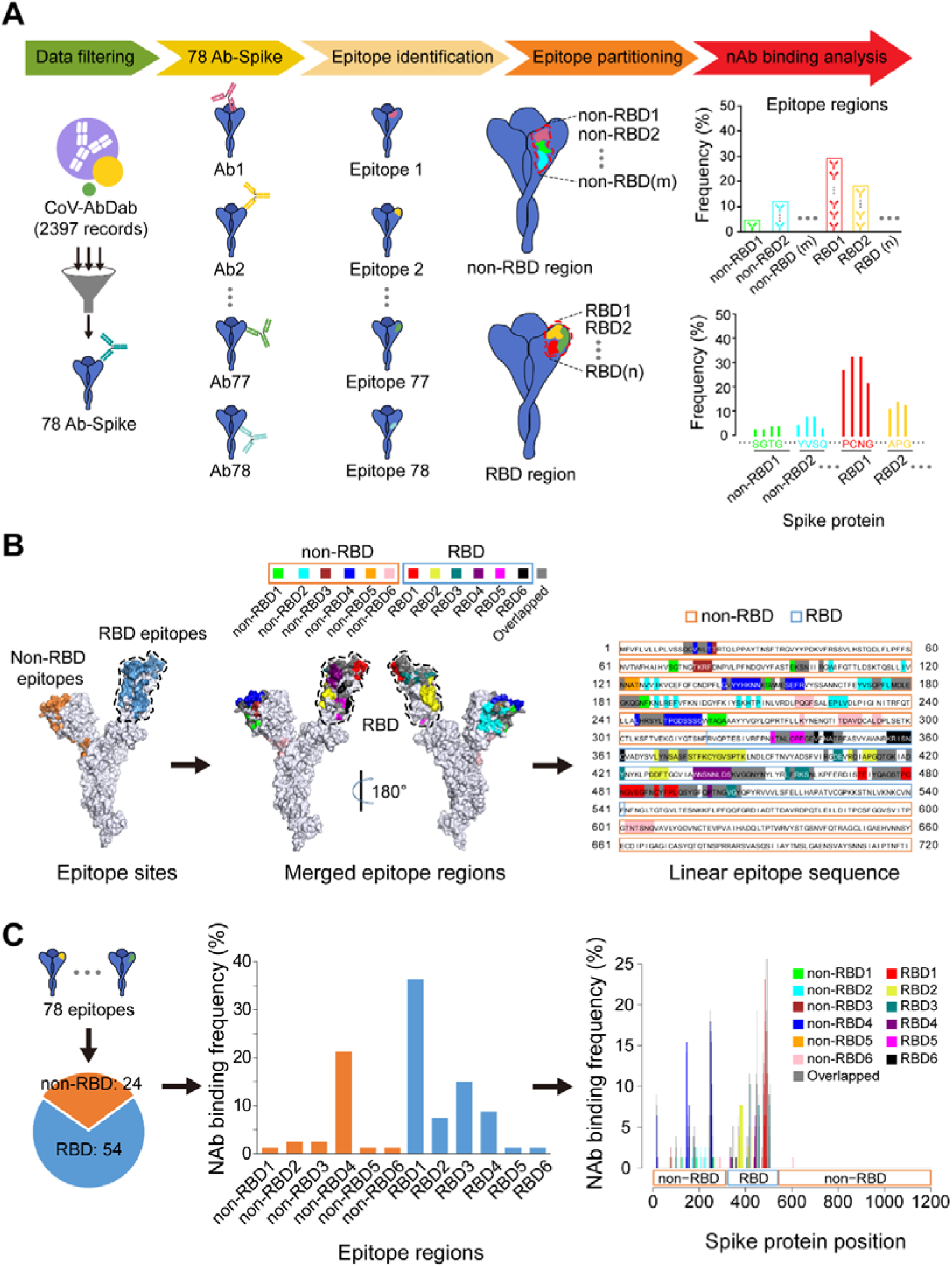
Non-RBD and RBD epitopes are identified within the spike protein of SARS-CoV-2. (A) Non-RBD and RBD epitope identification flow chart. (B) Left, identified non-RBD and RBD epitopes. Middle, non-RBD, and RBD epitope regions are projected onto protein 3D structures. Right, linear sequence of 12 epitope regions. (C) Left, the ratio of non-RBD and RBD epitope sites. Middle, nAb binding frequency of the non-RBD and RBD epitope regions. Right, Ab binding frequency of epitope residues within the different epitope regions. The gray vertical line represents epitope residues overlapped by at least two epitope regions.

We developed a new software tool called IBEIP (intelligent batch epitope identification and partitioning) for intelligent analysis to screen epitope sites and partition S antigen regions (Methods, Identifying and partitioning epitopes of neutralizing antibody). In contrast to the traditional epitope-partition method that relied on manual processing to partition epitope regions [36], our IBEIP can utilize an innovative iterative algorithm to intelligently partition hundreds of epitope sites at a minus-scale processing speed. In the current study, we defined the epitope site as the antigenic residues that are contacted (distance < 5 Å) and buried (buried surface area > 0 Å^2^) by neutralizing antibodies (nAbs) *via* the interaction of at least one hydrogen bond in the S-nAbs.

By running IBEIP software, a total of 78 S epitope sites were identified for the 78 S-nAbs. These S epitope sites were automatically partitioned into six non-RBD epitope regions (non-RBD1 to non-RBD6) and six RBD epitope regions (RBD1to RBD6) by IBEIP (Figure 3 (B and C), Supplemental Figure 3 and 4). It is noteworthy that five of the six non-RBD regions (non-RBD1 to non-RBD5) were located in the NTD of S protein and one spanned from NTD (residues 217-220, 278, 286-290, 294) to S1/S2 (residues 602-607) domain of the S protein (Figure 3 (B)). Furthermore, in the six non-RBD regions we identified a total of 24 epitope sites (non-RBD1: 1 site, non-RBD2: 2 sites, non-RBD3: 2 sites, non-RBD4: 17 sites, non-RBD5: 1 site, non-RBD6: 1 site) (Table 1), 54 epitope sites were identified in the six RBD regions (RBD1: 27 sites, RBD2: 6 sites, RBD3: 12 sites, RBD4: 7 sites, RBD5: 1 site, RBD6: 1 site) (Table 1).

**Figure 4.**
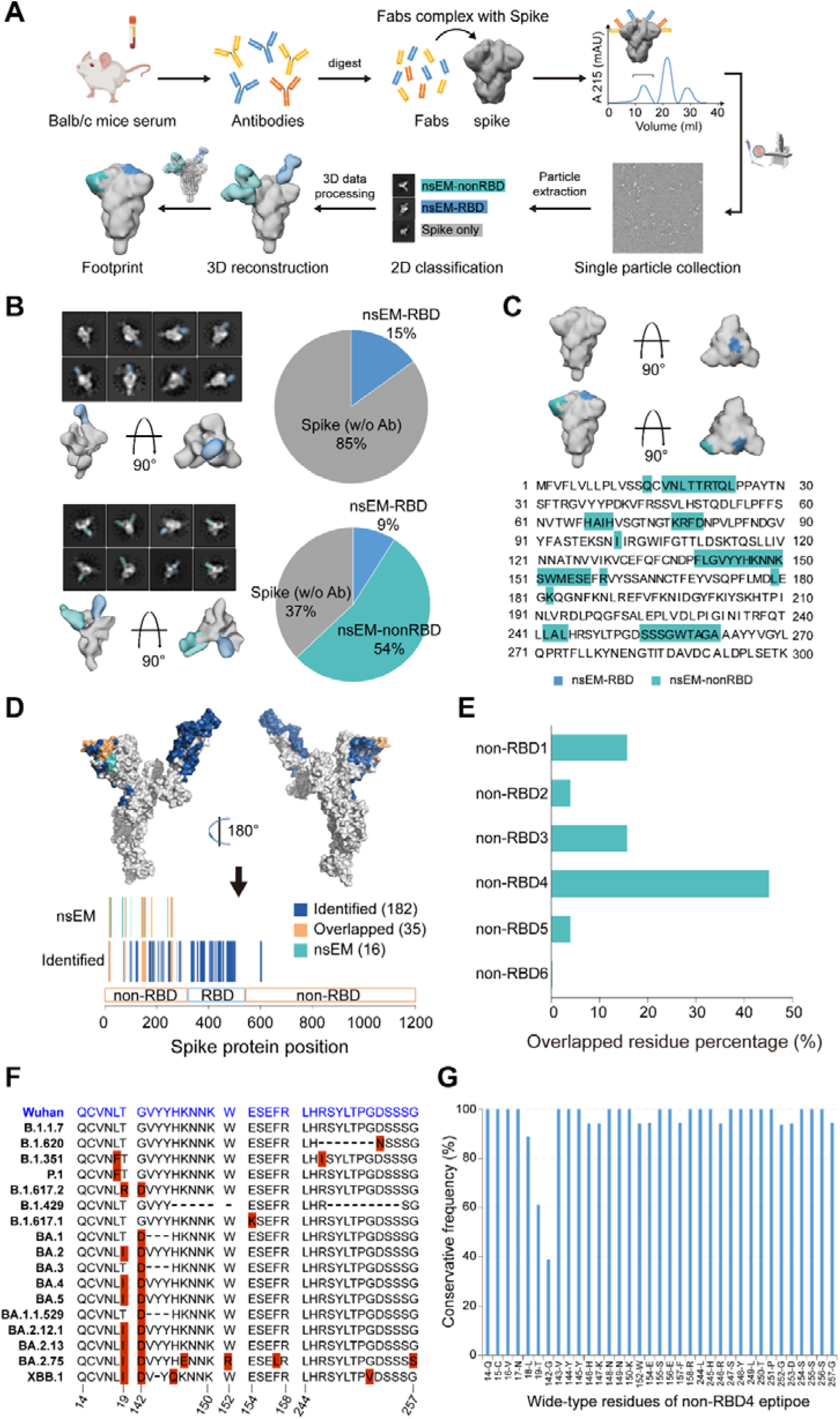
Ns-EMPEM analysis of polyclonal Fabs from vaccinated mouse sera complexed with SARS-CoV-2 spikes. (A) Schematic diagram of ns-EM polyclonal epitope mapping (EMPEM) analysis. (B) Representative 2D classes, side and top views of the 3D reconstructions from ns-EMPEM analysis of SARS-CoV-2 spikes complexed with polyclonal Fabs isolated from DNA(S)-RBD-RBD and DNA(S)-S-S vaccination groups. Pie charts show the proportion of Abs targeting the SARS-CoV-2 nsEM-nonRBD (sea green), nsEM-RBD (blue), and spike (w/o Ab) (gray). (C) Surface representation of each antibody epitope on the SARS-CoV-2 spike. The spike trimer with three “down” RBDs is adapted from PDB ID 6XR8. The epitope residues involved in NTD binding are labeled with their respective color. Interaction residues in NTD are defined by a 5 Å distance cut-off. (D) Comparison between identified and ns-EMPEM epitopes. (E) The percentage of ns-EMPEM non-RBD epitopes overlapped with the identified non-RBD epitope regions. (F) Residue sequences of the non-RBD4 epitope region in 17 SARS-CoV-2 variants. Mutated residues are highlighted with a red background and residue deletions are indicated by “-”. (G) Conservative frequency of the non-RBD4 epitope residues across 17 SARS-CoV-2 variants.

**Table 1:** The epitope residues and regions within the non-RBD and RBD regions on the spike protein of SARS-CoV-2.

| Antigen | Epitope regions | Ab binding frequency | Overlapped site positions | Unique site positions |
| --- | --- | --- | --- | --- |
| RBD | RBD1 | 34.62% | 403, 416, 417, 444-450, 456, 473-478, 486, 487, 489, 493-496, 498, 505 | 470, 471, 479-485, 488, 490-492 |
|  | RBD2 | 7.69% | 369, 370, 373, 374, 409, 414-417, 420, 460 | 368, 371, 372, 375-386, 408, 411-413, 427-430 |
|  | RBD3 | 15.38% | 369, 370, 403, 409, 414-417, 420, 449, 456, 460, 473-478, 486, 487, 489, 493-496, 498, 500-502, 505 | 405, 406, 421, 455, 457-459, 503, 504 |
|  | RBD4 | 8.97% | 345, 346, 373, 374, 444-450, 498, 500-502 | 436-443, 499 |
|  | RBD5 | 1.28% | 333-335, 339, 340 | 332, 336-338 |
|  | RBD6 | 1.28% | 333-335, 339, 340, 345, 346 | 341, 343, 344, 356-361 |
| Non-RBD | non-RBD1 | 1.28% | 97-99, 150, 152, 178-184, 245-249 | 71, 72, 96, 151, 185, 186, 259-262 |
|  | non-RBD2 | 2.56% | 99, 102, 121, 173, 174, 177-180, 182 | 104, 119, 126, 128, 170-172, 175, 176, 188, 190, 192, 205, 207, 209, 224-227 |
|  | non-RBD3 | 2.56% | 14, 15, 17, 18, 143, 247 | 20, 76-79 |
|  | non-RBD4 | 21.79% | 14, 15, 17, 18, 143, 150, 152, 154, 245-249 | 16, 19, 142, 144-149, 155-158, 244, 250-257 |
|  | non-RBD5 | 1.28% | 97-99, 102, 121, 154, 173, 174, 177, 178, 180-184 | 122-124 |
|  | non-RBD6 | 1.28% | / | 217-220, 278, 286-290, 293, 294, 602-607 |

We next located the important epitope regions by analyzing the nAb binding frequency among these epitope regions. The binding frequency is defined as the percentage of nAbs that bind to the specific epitope region. We found that the nAb binding frequencies of non-RBD4 and RBD1 were significantly higher than those of other regions, 21.79% and 34.62%, respectively (Figure 3 (C) and Table 1). In addition, the nAb binding frequency also showed significant differences among epitope residues within the same epitope region (Figure 3 (C)). The top 10 frequency residues of non-RBD4 were residues 144YYHKN148 (14.1%-15.38%) and 250TPGDS254 (10.26%-17.95%). The top 7 frequency residues of RBD1 were residues 481NGVEG485 (10.25%-23.08%), 490F (11.54%), and 492L (16.67%). More importantly, we observed a significant overlap between the non-RBD4 region (residues 14-19, 142-150, 152, 154-158 and 244-257) identified herein and a previously reported NTD supersite epitope region (residues 14–20,140–158 and 245–264) [13,37]. This overlap underscores the reliability of our antigenic epitope identification and partitioning methodology. These results indicated that the non-RBD4 region might be important for the conformational non-RBD domain to induce the neutralizing antibody response against the SARS-CoV-2 virus.

### Quantifying the percentage of conformational epitope-specific binding antibodies

To further validate the importance of the above-identified epitope region for enhancing cross-neutralization in mice with conformational non-RBD region boost, we employed negative stain electron microscopy-based polyclonal epitope mapping (ns-EMPEM) to elucidate the epitopes of serum antibodies that bind to the conformational non-RBD and RBD region of SARS-CoV-2 spike protein (Figure 4 (A)). Polyclonal Fabs were obtained by papain digested IgGs pooled from S_DNA_-RBD-RBD/S_DNA_-S-S group sera. Subsequently, the resulting Fabs were complexed with the spike protein and purified using size exclusion chromatography for single-particle EM data collection. Standard EM processing was performed [38], including particle extraction, 2D classification, and 3D reconstruction, while extensive 3D classification was conducted to sort each Fab specificity. The spike model (PDB: 6XR8) and Fab model (PDB: 6WS6) were subsequently fitted into reconstructed 3D maps to identify epitopes of each antibody on the surface of the SARS-CoV-2 spike protein. The conformation of antigen S protein manifested a prototypical homotrimer structure (Supplemental Figure 5). The 3D reconstruction of polyclonal Fabs in complex with the wild-type spike protein of SARS-CoV-2 from the S_DNA_-RBD-RBD group (Figure 4 (B), left and up) and the S_DNA_-S-S group (Figure 4 (B), left and down) revealed significant differences in their binding specificities. The binding responses of polyclonal Fabs to RBD were observed in the S_DNA_-RBD-RBD group, whereas the S_DNA_-S-S group antibodies exhibited binding to both conformational non-RBD and RBD epitopes. Moreover, our findings indicate that non-RBD-targeting antibodies approach spike protein from two distinct angles, which were labeled as ‘nsEM-nonRBD1’ and ‘nsEM-nonRBD2’ (Supplemental Figure 6). In the S_DNA_-RBD-RBD group, RBD antibodies accounted for approximately 15% of the total ∼13k particles (Figure 4 (B), right and up). In contrast, in the S_DNA_-S-S group, non-RBD antibodies accounted for approximately 54% of the total ∼10k particles, while RBD antibodies occupied only 9% of the total particles (Figure 4 (B), right and down). It can be seen that among all the antibodies induced by the S protein, there are significantly more antibodies targeting the non-RBD region than those targeting the RBD region. Building upon the above epitope region analysis (Figure 4 (B and C)), most non-RBD epitope regions are located in the NTD region. We believe the significant advantage of cross-neutralization potency in the S_DNA_-S-S group over the S_DNA_-RBD-RBD group (Figure 2 (F)) is attributable to the low mutational variability and the relative epitope conservation on the NTD region. Additionally, a list of potential contact residues between the spike protein and non-RBD antibodies was generated with an interaction distance of ≤5Å (Figure 4 (C), right). However, reconstructing the RBD antibodies was more challenging, and it was not possible to identify concrete interaction residues due to the flexible nature of the RBD [38]. Nevertheless, the analysis of EM data (2D and 3D) revealed the presence of both non-RBD and RBD antibodies.

In this study, the epitope profile identified by ns-EMPEM analysis was observed in accordance with the identified epitopes of the SARS-CoV-2 S protein (Figure 4 (D)). By in-depth analyses of the distribution of the non-RBD epitope sites identified by ns-EMPEM within the 6 identified non-RBD epitope regions (non-RBD1 to non-RBD6), we observed that the residues of nsEM-nonRBD overlapped 45.1% with the non-RBD4 epitope region (Figure 4 (E)). The majority of epitope residues (32 out of 35) in the non-RBD4 region were highly conserved (conserved frequency > 93 %) across 17 SARS-CoV-2 variants, including 13 variants of concern and 3 variants of interest (Figure 4 (F and G)). Overall, the non-RBD epitopes identified by ns-EMPEM were consistent with our identified non-RBD4 epitope region and validated its importance for enhanced cross-neutralization in mice with conformational non-RBD region boost.

### Validating the contribution of non-RBD antibodies to neutralization

A pseudotyped virus neutralization assay was conducted to validate the impact of NTD-specific antibodies on the neutralizing efficacy of the serum (Figure 5 (A)). The neutralization potency of serum from the S_DNA_-S-S group decreased upon blocking with a low concentration of NTD protein (0.04 μg/μL) (Figure 5 (B, D)). Furthermore, an elevation in the blocking concentration of NTD protein (0.08 μg/μL) led to a significant decrease in serum neutralization potency (*P* < 0.005, Figure 5 (C, D)). These findings provide evidence that NTD-specific antibodies play a crucial role in enhancing the neutralization efficacy against SARS-CoV-2.

**Figure 5.**
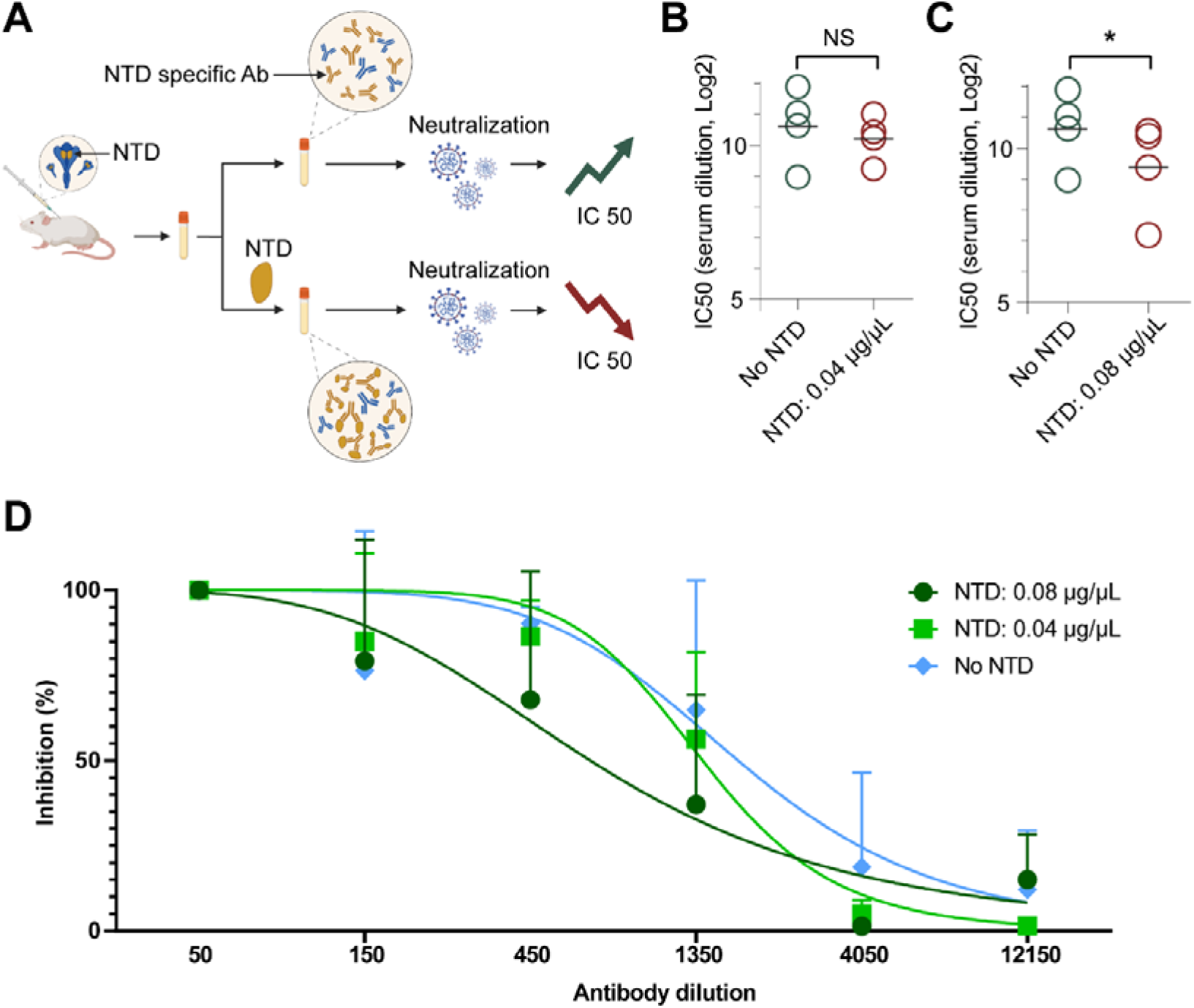
Contribution of non-RBD antibodies to neutralization. (A) Schematic diagram of the experiment to evaluate the influence of NTD-specific Abs on neutralization against pseudotyped virus. (B-C) Pseudotyped virus neutralization titers of serum with or without NTD protein incubation. (D) Fitting analysis of pseudotyped virus neutralization titers of serum with different NTD protein concentrations. \**P* < 0.05 (two-tailed Student t-test). NS, not significant.

## Discussion

Enhancing the broad-spectrum efficacy of vaccines is crucial for controlling the spread of highly variable pathogens like the SARS-CoV-2 virus. Multiple hotspot mutations in the RBD region of spike antigen resulted in enhanced immune evasion for SARS-CoV-2 viruses [39–41]. Furthermore, the therapeutic effect of neutralizing antibodies targeting the RBD region significantly reduced for SARS-CoV-2 variants [1,8]. In contrast, the non-RBD region of the spike protein shows fewer immune evasion mutations and antibodies targeting this region exhibit good broad-spectrum neutralization against SARS-CoV-2 viruses [11–13]. Therefore, we focus on the non-RBD epitopes of the spike protein to enhance broad-spectrum efficacy for SARS-CoV-2 vaccines. In this study, by using the ‘DNA prime-protein boost’ strategy in a mouse model, we demonstrated that the high ratio of antibodies targeting conformational non-RBD region can significantly enhance cross-neutralization against SARS-CoV-2 viruses. By combining intelligentized epitope identification and electron microscopy (EM) analysis, we proved that the high ratio of binding antibodies targeting NTD supersite in the non-RBD region contributed to the broad cross-neutralization against SARS-CoV-2 viruses.

The maintenance of antigen spatial conformation is closely related to the quality of vaccine-induced antibody responses. Currently, for the construction of SARS-CoV-2 vaccines, the preservation of the spatial conformation of either full-length S or RBD antigens has been realized [21,42,43]. In contrast, conformational non-RBD antigens have still not been successfully developed yet, although several studies have highlighted the importance of conformational non-RBD antigens in fighting against SARS-CoV-2 infection [26,44]. One of the challenges in developing conformational non-RBD antigens is that *in vitro* expression of non-RBD will significantly alter its spatial conformation [28]. This challenge requires us not to directly inject protein antigen to boost conformational non-RBD-specific antibodies. To solve this problem, we used full-length S antigen to elicit an immune response against both non-RBD and RBD regions and compared it with an immune response against RBD antigen to assess the potential of non-RBD region in broad cross-neutralization against SARS-CoV-2 viruses.

DNA prime-protein boost strategy has been proven to be a robust immunization approach in our and other groups’ vaccine models such as SARS-CoV-2, HIV, and HBV [33,45,46]. Therefore, DNA prime-protein boost can provide a reasonable strategy for inducing potent conformational non-RBD-specific antibody responses. In our study, these non-RBD boost mice displayed a higher neutralization potency (from 3.5 to 8.24 fold) against diverse SARS-CoV-2 viruses (Wuhan-Hu-01, B.1.617.2, and B.1.1.529), in comparison to the mice without non-RBD boost. These results demonstrated that conformational non-RBD antigens play a more important role in improving the broad cross-neutralization of the SARS-CoV-2 vaccine.

Another novelty of our study is the combination of intelligent epitope identification with electron microscopy-based (ns-EMPEM) epitope validation to quickly reveal the underlying mechanism for enhanced cross-neutralization capacity in non-RBD boost mice. To identify potentially crucial epitopes that are responsible for enhancing the cross-neutralizing capability of S-boosted animals, we developed a new intelligent iteration method (IBEIP) to analyze the electron microscope structures of SARS-CoV-2 virus bound to nAbs from the CoV-AbDab database and identify their corresponding epitope sites and regions. By analyzing the antibody binding frequency against these epitope sites, we identified two significant epitope regions, namely non-RBD4 and RBD1. The non-RBD4 region mainly overlapped with the NTD supersite. The ns-EMPEM analysis further pinpointed the pivotal epitope positions and quantified the percentage of antibodies targeting the conformational non-RBD region. The non-RBD boost mice with potent SARS-CoV-2 cross-neutralization obtained 54% non-RBD-specific antibody responses and 9% RBD-specific antibody responses. The high proportion of NTD-specific antibody responses may be linked to the effects of DNA priming, which, as shown in previous studies, can induce T follicular helper (Tfh) cell responses, promote germinal center (GC) B cell formation, and enhance antigen-specific B cell responses [46–48]. Notably, the ns-EMPEM analysis also indicated that non-RBD-specific Abs predominantly focused their response on the non-RBD4 region (NTD supersite) which further affirms the accuracy of our developed IBEIP approach.

Furthermore, *via* blocking the NTD-specific binding antibodies, the neutralization potency of serum samples from non-RBD boost mice significantly decreased. Such a decrease in neutralization potency has a close relationship with blocking NTD-specific binding antibodies. Therefore, these results indicated that the high ratio of neutralizing antibodies targeting the NTD supersite plays a crucial role in mediating SARS-CoV-2 cross-neutralization.

In summary, this work represents a proof-of-concept investigation of the pivotal role of conformational non-RBD epitopes in enhancing the broad cross-neutralization of SARS-CoV-2 vaccines. Our findings offer a fresh research perspective and a design strategy for next-generation SARS-CoV-2 vaccines. Our approach of combining intelligent epitope identification with electron microscopy-based epitope validation enabled a rapid and accurate determination of critical epitope information for the enhanced broad cross-neutralizing response of conformational non-RBD region. Such an approach would also be beneficial to the prevention and treatment research of other pathogens that have abundant crystal structures of antigen-antibody complexes, such as HIV or influenza. To further validate the broad applicability of our research strategy, additional studies in non-human primate models are needed. Non-human primates share closer physiological and immune system similarities to humans than mice, which may provide more relevant insights into the translational potential of our approach.

## Funding

This work was supported by the National Key R&D Program of China (2022YFA1206400); the National Natural Science Foundation of China (22277017); the Strategic Priority Research Program of Chinese Academy of Sciences (XDB0770000); the Science and Technology Plan-biological medical special project of Yunnan Province (202302AA310005); the Basic Research Program of Yunnan Province (202201AU070152, 202101AS070051, 202002AA100009, 202101AU070126); Leading Medical Talents Program of Health Commission of Yunnan Province (L-2018013); the Young and Middle Aged Academic and Technical Leaders Program of Yunnan Province (202005AC160010); the National Natural Science Foundation of China (82304307); the National Basic Research Program of China (2020YFA0710700); the Fundamental Research Funds for the Central Universities (3332022073).

## Author’s contributions

Y. Liu, Y. R. Yang, and C. Chen led the project, designed the experiments, interpreted the data, and drafted the manuscript. Y. F. Ma implemented the Python code, analyzed and interpreted data from PDB and CoV-AbDab, and drafted the manuscript. Y. F. Ma, B. Xie, K. Chen, J. Zhu, and X. He carried out the experiments and data analysis. All authors discussed the results and commented on the manuscript.

## Disclosure statement

No potential conflict of interest was reported by the author(s).

---

**Supplemental Figure 1.**
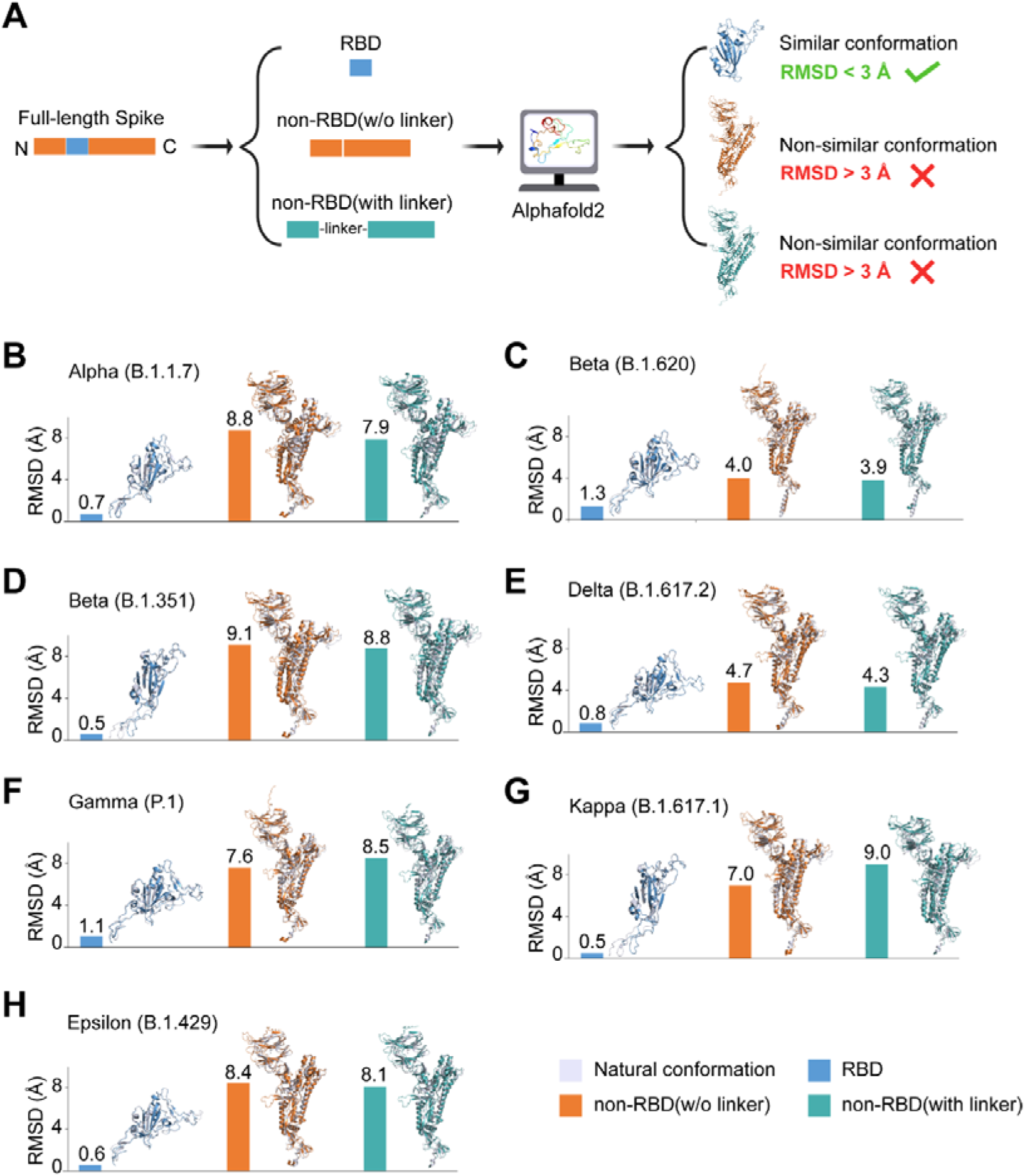
Structure analysis of conformational non-RBD and RBD subunits. (A) Selection strategy of conformational non-RBD and RBD protein vaccines. (B-H) Structural comparison of predicted non-RBD and RBD epitope vaccines with natural conformation of full-length S antigen.

**Supplemental Figure 2.**
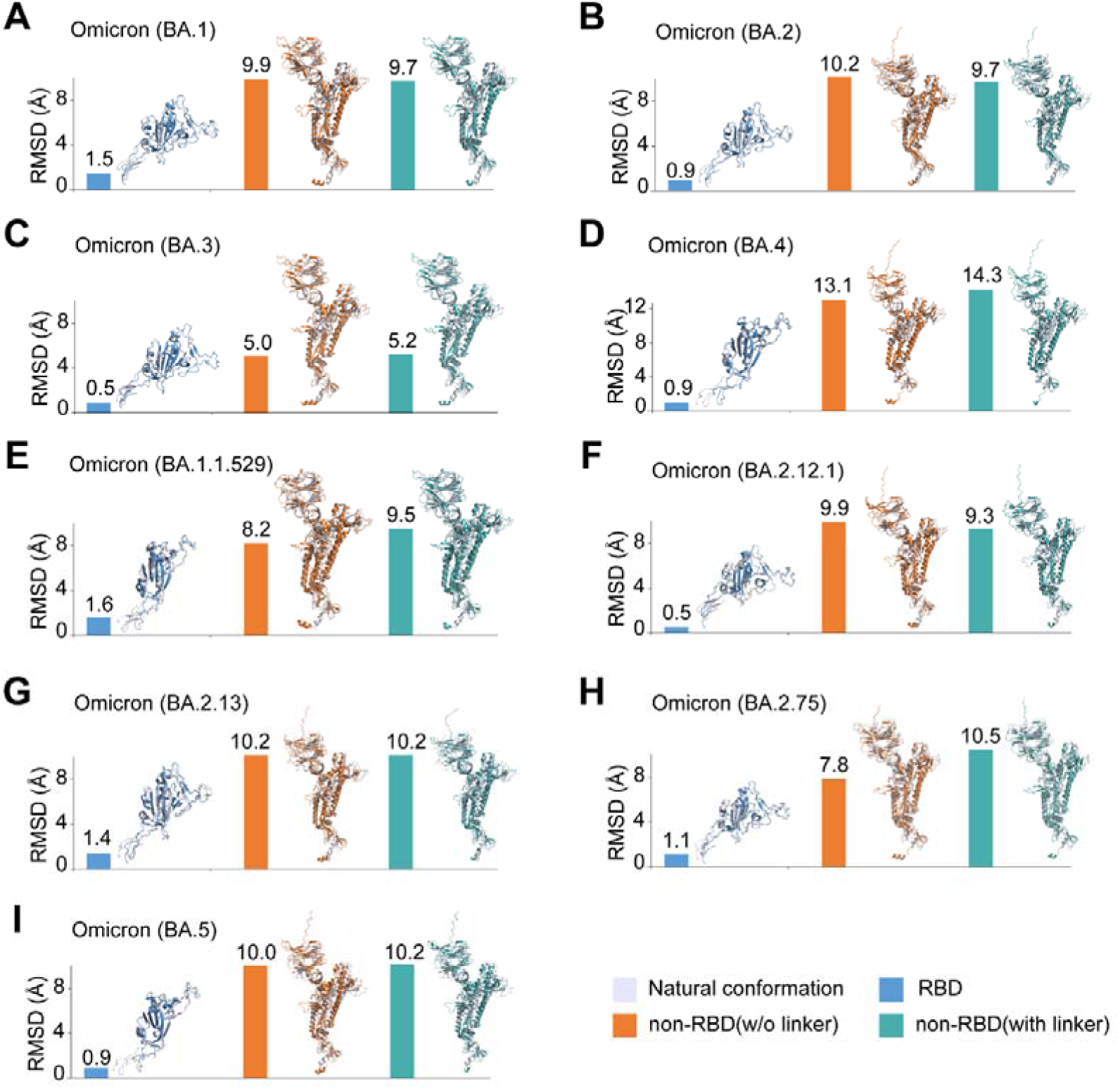
Structure comparison of predicted non-RBD and RBD epitope vaccines with spatial conformation of full-length S antigen for SARS-CoV-2 Omicron variants.

**Supplemental Figure 3.**
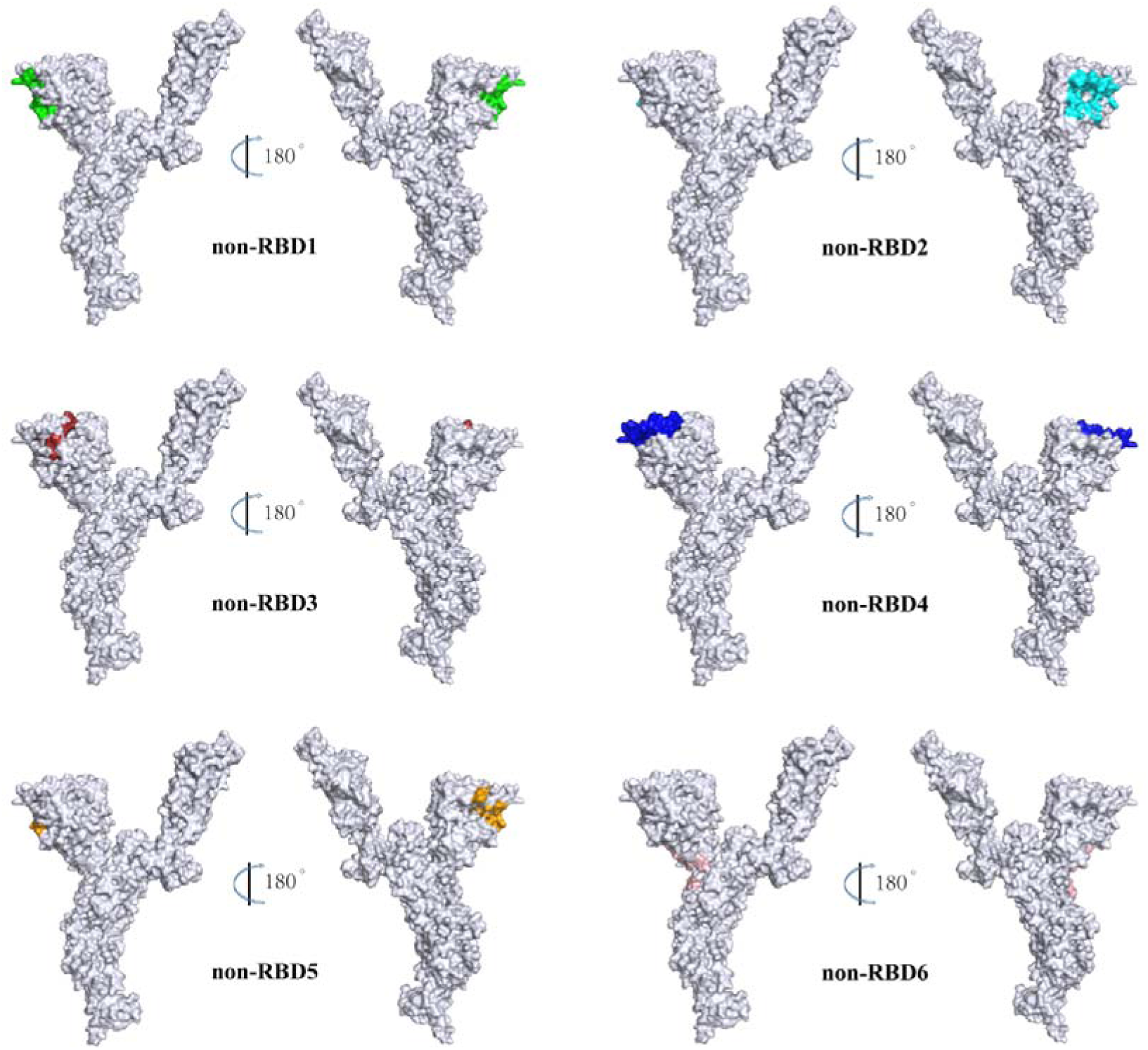
The six identified regions of non-RBD epitope.

**Supplemental Figure 4.**
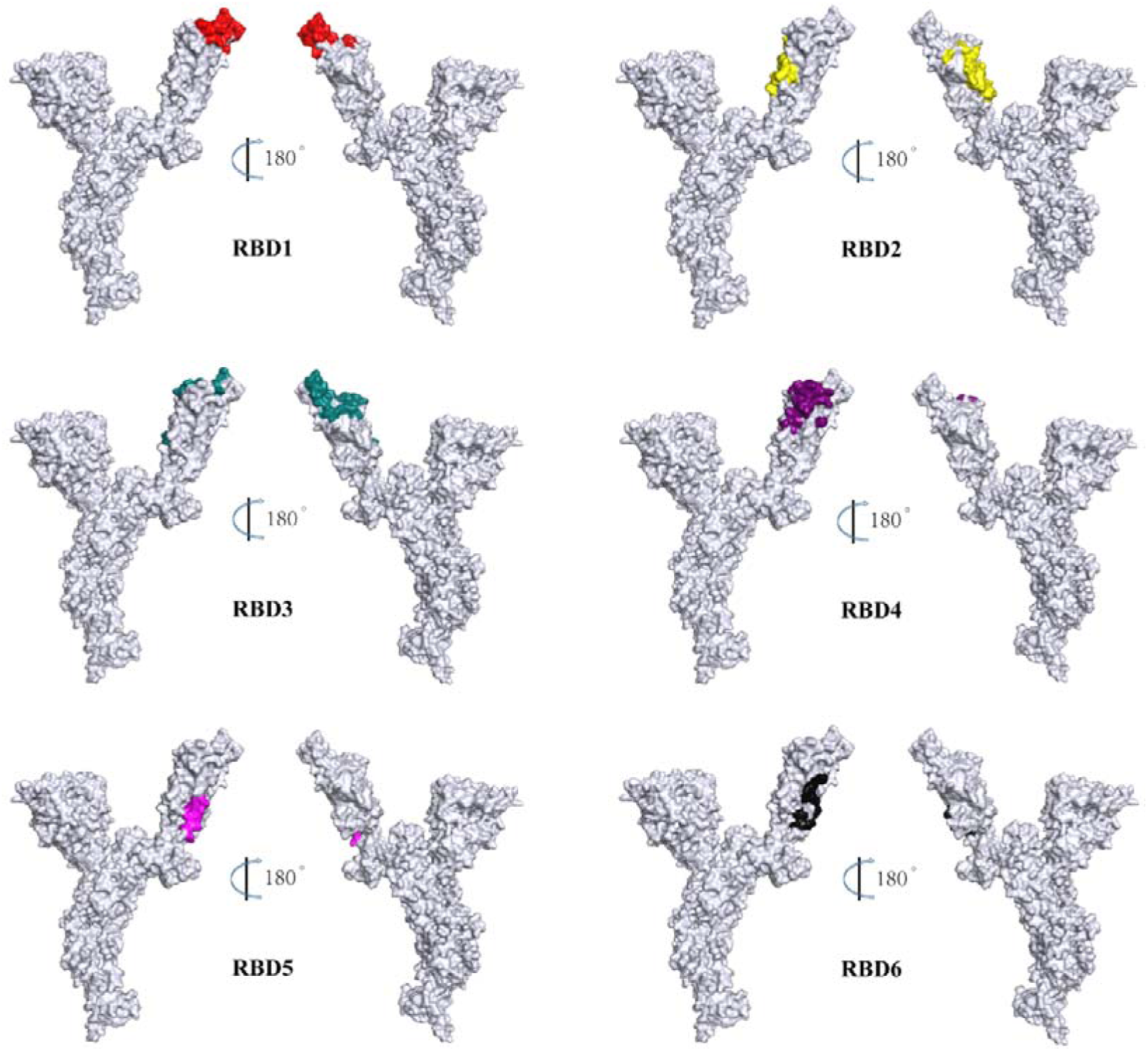
The six identified regions of RBD epitope.

**Supplemental Figure 5.**
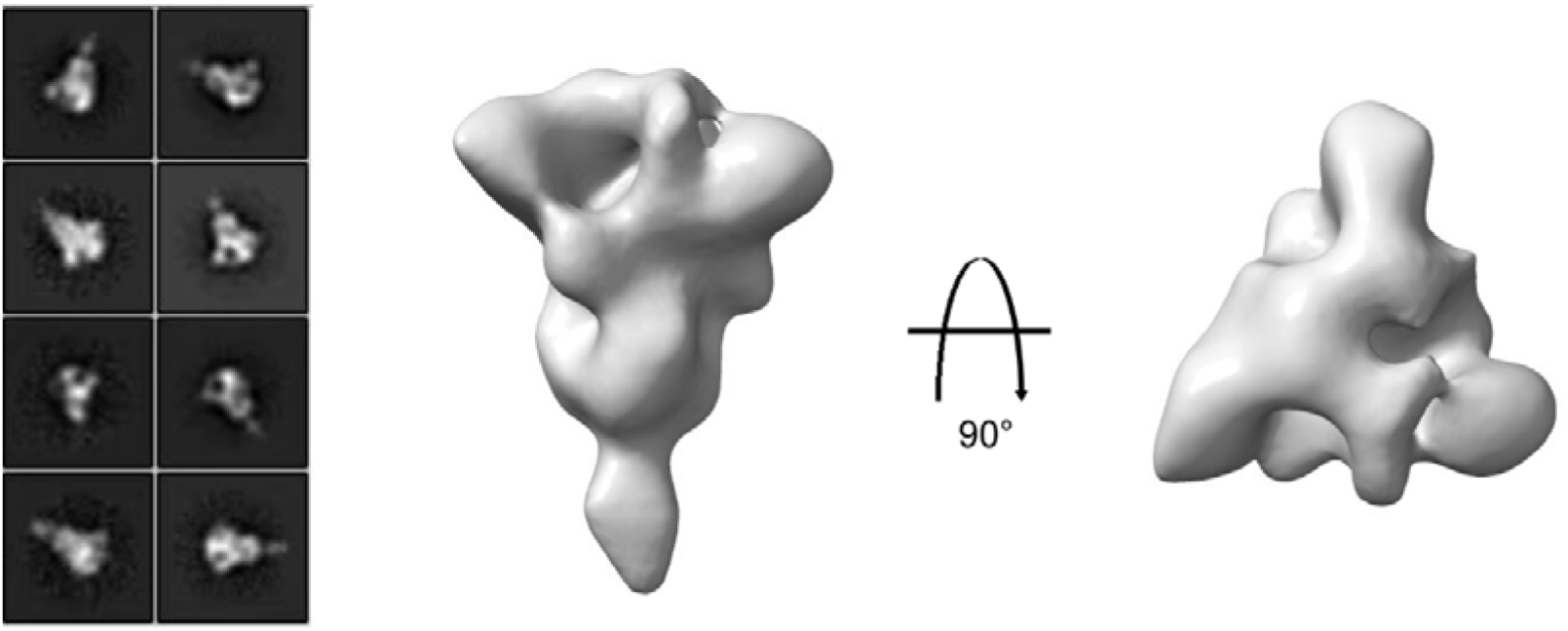
The 3D reconstructions from ns-EMPEM analysis of SARS-CoV-2 spike protein.

**Supplemental Figure 6.**
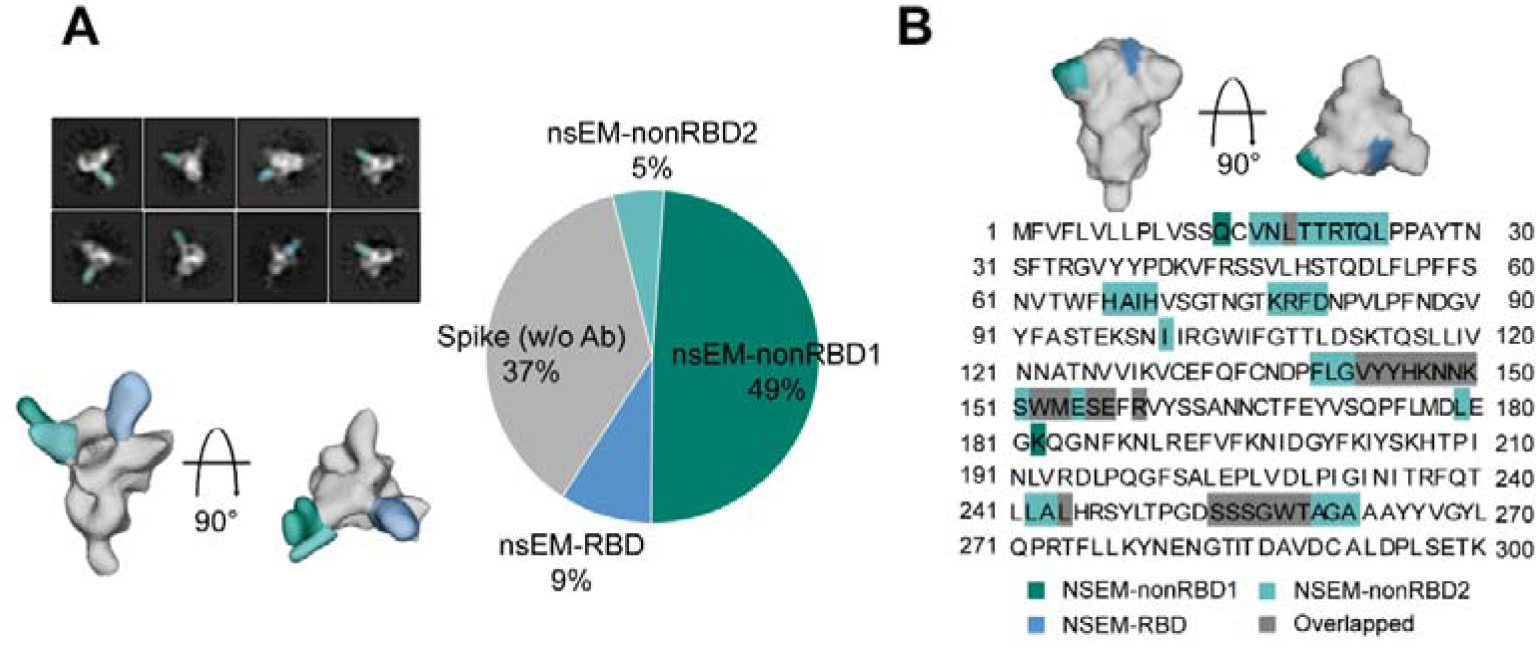
The ns-EMPEM analysis revealed two non-RBD epitope regions. **(A)** Representative 2D classes, side and top views of the 3D reconstructions from ns-EMPEM analysis of SARS-CoV-2 spikes complexed with polyclonal Fabs isolated from DNA(S)-RBD-RBD and DNA(S)-S-S vaccination groups. Pie charts show the proportion of Abs targeting the SARS-CoV-2 nsEM-nonRBD1 (teal), nsEM-nonRBD2 (cerulean), nsEM-RBD (blue), and spike (w/o Ab) (gray). **(B)** Surface representation of each antibody epitope on the SARS-CoV-2 spike. The spike trimer with three “down” RBDs is adapted from PDB ID 6XR8. The epitope residues involved in NTD binding are labeled with their respective color. Interaction residues in NTD are defined by a 5 Å distance cut-off.

## Notes

### Competing Interest Statement

The authors have declared no competing interest.

### Summary of Updates

We thoroughly reviewed the original manuscript and corrected several typographical and grammatical errors. We carefully revised the main text to enhance clarity and readability. All figures in the manuscript have been updated for improved clarity and accuracy.

