## Supplemental Table 1 for "Enhanced Antibody Response to the Conformational Non-RBD Region *via* DNA Prime-Protein Boost Elicits Broad Cross-Neutralization Against SARS-CoV-2 Variants"

| **SN** | **WHO name** | **Strain** | **RMSD (Å)** | | | **PDB ID** | **References** |
| --- | --- | --- | --- | --- | --- | --- | --- |
|  |  |  | **RBD** | **non-RBD(w/o linker)** | **non-RBD(with linker)** |  |  |
| 1 | WildType | Wuhan | 1.4 | 7.4 | 7.2 | 7NTC | [1] |
| 2 | Alpha | B.1.1.7 | 0.7 | 8.8 | 7.9 | 8DLJ | [2] |
| 3 | Beta | B.1.620 | 1.3 | 4.0 | 3.9 | 7YBK | To be published |
| 4 | Beta | B.1.351 | 0.5 | 9.1 | 8.8 | 8DLM | [2] |
| 5 | Gamma | P.1 | 1.1 | 7.6 | 8.5 | 8DLO | [2] |
| 6 | Delta | B.1.617.2 | 0.8 | 4.7 | 4.3 | 7Y6D | [3] |
| 7 | Epsilon | B.1.429 | 0.6 | 8.4 | 8.1 | 8DLX | [2] |
| 8 | Kappa | B.1.617.1 | 0.5 | 7.0 | 9.0 | 7TF0 | [4] |
| 9 | Omicron | BA.1 | 1.5 | 9.9 | 9.7 | 8HFX | [5] |
| 10 | Omicron | BA.2 | 0.9 | 10.2 | 9.7 | 7XIX | [6] |
| 11 | Omicron | BA.3 | 0.5 | 5.0 | 5.2 | 7Y20 | [7] |
| 12 | Omicron | BA.4 | 0.9 | 13.1 | 14.3 | 8H07 | [8] |
| 13 | Omicron | BA.5 | 0.9 | 10.0 | 10.2 | 8GTQ | [9] |
| 14 | Omicron | BA.1.1.529 | 1.6 | 8.2 | 9.5 | 8DZI | [10] |
| 15 | Omicron | BA.2.12.1 | 0.5 | 9.9 | 9.3 | 8CIM | To be published |
| 16 | Omicron | BA.2.13 | 1.4 | 10.2 | 10.2 | 7XNR | [6] |
| 17 | Omicron | BA.2.75 | 1.1 | 7.8 | 10.5 | 8GS6 | [11] |
| 18 | Omicron | XBB.1 | 1.0 | 8.0 | 8.0 | 8IOU | [12] |

**Supplementary Table 1.** **Structure comparison of non-RBD and RBD subunits within different SARS-CoV-2 strains.**
